# African nonhuman primates are infected with the yaws bacterium *Treponema pallidum* subsp. *pertenue*

**DOI:** 10.1101/135491

**Authors:** Sascha Knauf, Jan F. Gogarten, Verena J. Schuenemann, Hélène M. De Nys, Ariane Düx, Michal Strouhal, Lenka Mikalová, Kirsten I. Bos, Roy Armstrong, Emmanuel K. Batamuzi, Idrissa S. Chuma, Bernard Davoust, Georges Diatta, Robert D. Fyumagwa, Reuben R. Kazwala, Julius D. Keyyu, Inyasi A. V. Lejora, Anthony Levasseur, Hsi Liu, Michael A. Mayhew, Oleg Mediannikov, Didier Raoult, Roman M. Wittig, Christian Roos, Fabian H. Leendertz, David Šmajs, Kay Nieselt, Johannes Krause, Sébastien Calvignac-Spencer

**Author notes:** Correspondence to: Dr. Sebastien Calvignac-Spencer, Robert Koch Institute, Seestrasse 10, 13353 Berlin, Germany; Prof. David Šmajs, Faculty of Medicine, Masaryk University, 625 00 Brno, Czech Republic; Prof. Dr. rer. nat. Johannes Krause, Max Planck Institute for the Science of Human History, Jena, Germany. Current affiliation: Unité Mixte Internationale 233, Institut de Recherche pour le Développement, INSERM U1175, and University of Montpellier, Montpellier, France (Hélène M. De Nys). These authors contributed equally to this work (Sascha Knauf, Jan F. Gogarten, Verena J. Schuenemann, Hélène M. De Nys, David Šmajs, Kay Nieselt, Johannes Krause, Sébastien Calvignac-Spencer).

## Abstract

*Treponema pallidum* subsp. *pertenue* (*TPE*) is the causative agent of yaws. The disease was subject to global eradication efforts in the mid 20^th^ century but reemerged in West Africa, Southern Asia, and the Pacific region. Despite its importance for eradication, detailed data on possible nonhuman disease reservoirs are missing. A number of African nonhuman primates (NHPs) have been reported to show skin ulcerations suggestive of treponemal infection in humans. Furthermore antibodies against *Treponema pallidum* (*TP*) have been repeatedly detected in wild NHP populations. While genetic studies confirmed that NHPs are infected with *TP* strains, subspecies identification was only possible once for a strain isolated in 1966, pinpointing the involvement of *TPE*. We therefore collected a number of recently isolated simian *TP* strains and determined eight whole genome sequences using hybridization capture or long-range PCR combined with next-generation sequencing. These new genomes were compared with those of known human *TP* isolates. Our results show that naturally occurring simian *TP* strains circulating in three African NHP species all cluster with human *TPE* strains and show the same genomic structure as human *TPE* strains. These data indicate that humans are not the exclusive host for the yaws bacterium and that a One Health approach is required to achieve sustainable eradication of human yaws.

## Introduction

The bacterium *Treponema pallidum* (*TP*) causes human syphilis (subsp. *pallidum*; *TPA*), bejel (subsp. *endemicum*; *TEN*) and yaws (subsp. *pertenue*; *TPE*). While syphilis reached a world-wide distribution (1, 2), bejel and yaws are endemic diseases. Bejel is found in the dry areas of Sahelian Africa and Saudi Arabia, whereas yaws is present in the humid tropics (3). Yaws is currently reported endemic in 13 countries and an additional 76 countries have a known history of human yaws but lack recent epidemiological data (4). The disease was subject to global eradication efforts in the mid 20^th^ century but reemerged in West Africa, Southern Asia, and the Pacific region (5). New large-scale treatment options triggered the ongoing, second eradication campaign that aims to eradicate human yaws globally by 2020 (5). Despite its importance for eradication, detailed data on possible nonhuman disease reservoirs are missing, which unfortunately contributed to the perception that *TPE* is an exclusive human pathogen.

Reports of *TP* infections in nonhuman primates (NHPs), however, have accumulated (6). A number of African NHPs have been reported to show skin ulcerations suggestive of treponemal infection in humans and antibodies against *TP* have been repeatedly detected in wild NHP populations (7-10). Genetic studies confirmed that monkeys and great apes are infected with *TP* strains (11-13). However, most of these analyses only determined short DNA sequences and therefore only collected information on a small number of polymorphic sites considered to segregate between the *TP* subspecies (11, 13). Since sporadic recombination events between subspecies have been reported (14-16), a clear assignment of these strains to a *TP* subspecies was not possible. The only simian strain whose whole genome was sequenced - Fribourg-Blanc, isolated from a Guinea baboon (*Papio papio*) in 1966 (7) - unambiguously clustered with human *TPE* strains (GenBank: NC_021179.1) (17).

A fundamental question with regard to human yaws eradication is whether humans and NHPs are infected with the same pathogen. To address this question, we collected eight additional simian *TP* strains, determined their whole genome sequences using hybridization capture or long-range PCR combined with next-generation sequencing (NGS) and compared these new genomes with those of known human *TP* isolates (1, 12).

## Results and Discussion

### Humans are not the exclusive host for the yaws bacterium

We investigated three NHP species (*Cercocebus atys*, *Chlorocebus sabaeus*, and *Papio anubis*) from four independent populations in West and East Africa (**Table 1**). Samples were collected at Taï National Park (TaïNP; Côte d’Ivoire), Bijilo Forest Park (BFP, the Gambia), Niokolo-Koba National Park (NKNP, Senegal), as well as Lake Manyara National Park (LMNP, Tanzania). Monkeys either presented yaws-like orofacial and limb (TaïNP, BFP) or ulcerative anogenital skin lesions (BFP, NKNP, LMNP (18)). *TP* infection in olive baboons (*P. anubis*) at LMNP has already been confirmed (*8*). Via PCR, we were able to show the presence of *TP* in skin lesion biopsies or swabs from NHPs at TaïNP (*C. atys*) as well as BFP and NKNP (*C. sabaeus*).

**Table 1.** Non-human primates anesthetized for this study.

| Species | Number of individuals | Clinical manifestations | Park, country |
| --- | --- | --- | --- |
| Sooty mangabey (*Cercocebus atys*) | 5 | Face or lower extremities incl. bone deformation | TaïNP, Côte d'Ivoire |
| African green monkey (*Chlorocebus sabaeus*) | 5 | Face and anogenital | BFP, The Gambia |
| African green monkey (*Chlorocebus sabaeus*) | 3 | Genital | NKNP, Senegal |
| Olive baboon (*Papio anubis*) | 2 | Anogenital | LMNP, Tanzania |

The most suitable samples from all four populations were selected for whole genome sequencing. Selection criteria were based on high *TP* copy number or the ability to amplify long PCR fragments. To overcome the large background of host genomic DNA we used targeted DNA capture, coupled with NGS (1, 12, 19). Following quality filtering, removal of PCR duplicates, merging of different sequencing runs from the same sample, and mapping against the *TPE* strain Fribourg-Blanc (GenBank: NC_021179.1) reference genome, we obtained a range of 22,886-470,303 DNA sequencing reads per sample. Samples showing at least 80% coverage of the reference genome with depth coverage of three or higher were further analyzed. This resulted in a 6.1 to 121.0-fold average depth genome coverage for two samples per analyzed NHP population (n=8). For the reconstruction of a phylogenetic tree we included all available human *TP* strain genomes (1). Four potentially recombinant genes (*TPFB_0136*, *TPFB_0326*, *TPFB_0488*, *TPFB_0865*; (1)) were removed before a phylogenetic tree was constructed using the maximum parsimony method (**Fig. 1**).

**Figure 1.**
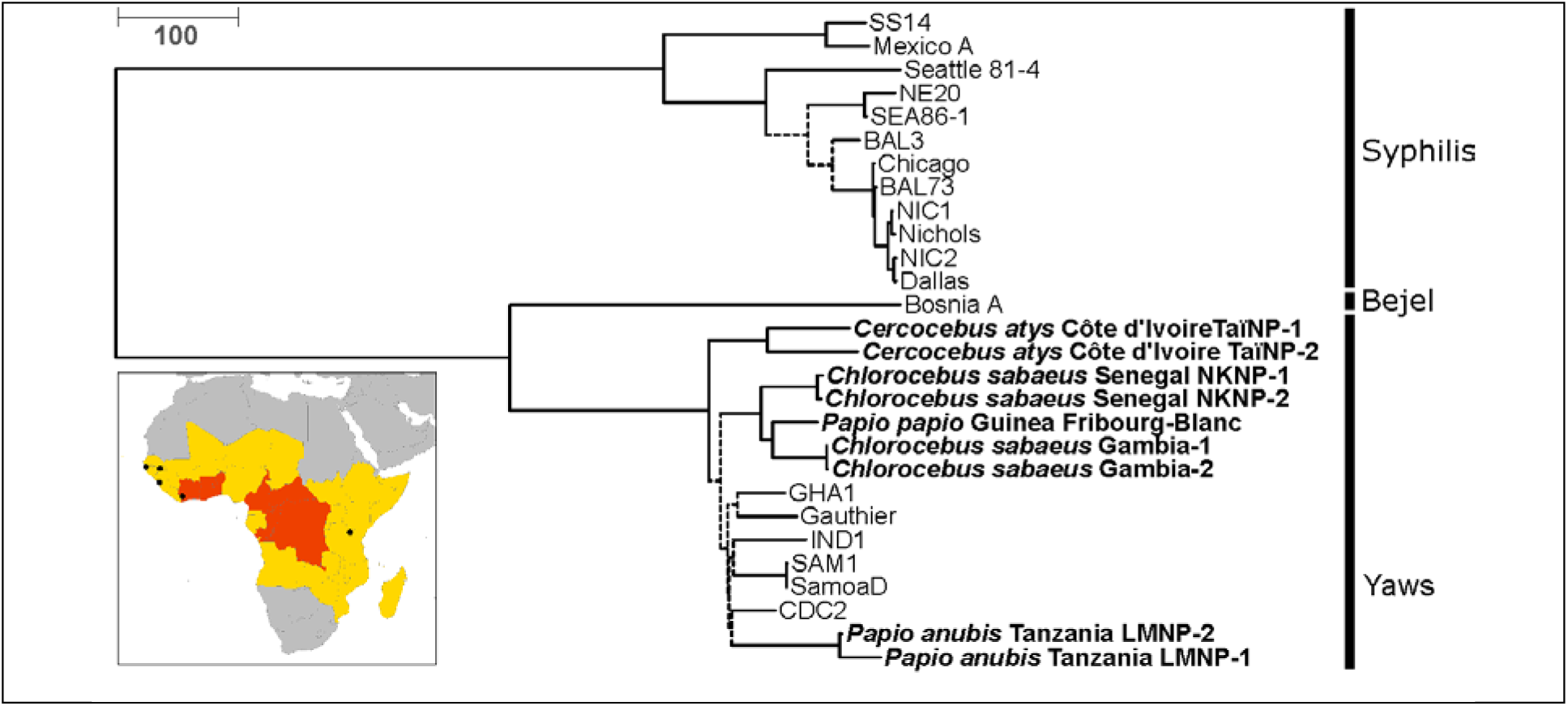
Phylogenomic analysis of NHP (in bold) and human-infecting *TP*. In this maximum parsimony tree, reconstructed using 2,063 informative positions, nodes that had less than 90% bootstrap support are indicated with dashed lines. Tip labels indicate the NHP species sampled, the country of origin, and the sample ID. Abbreviations used for human strains correspond to the classification in Arora et al. 2016 (1). The scale is in nucleotide substitution. Note that for this tree recombinant genes (*TPFB_0136*, *TPFB_0326*, *TPFB_0488*, *TPFB_0865*) were removed. The inset is a map of Africa where sites of origin of the NHP samples from which a *TP* genome was determined are indicated with black circles. A country’s 2013 yaws status based on the World Health Organization’s Global Health Observatory (http://www.who.int/gho/en/) is indicated by its color: grey indicates no previous history of yaws infections in humans, yellow indicates a country previously endemic for yaws though the current status is unknown, and countries in red indicate countries which are currently considered endemic for yaws.

Our results clearly show that naturally occurring simian *TP* strains cluster with human *TPE* strains, thus indicating that infection in African NHPs is caused by the *TP* subsp. *pertenue* (**Fig. 1**). Accordingly, humans are not the exclusive host for the yaws bacterium. Human yaws-causing *TPE* strains Samoa D (20), CDC-2 (21, 22), Ghana-051 [GHA1] (23), Kampung Dalan K363 [IND1] (1), and Gauthier (24) span a broad geographic and temporal range (at least four decades), but they are less divergent from each other than the two strains infecting sooty mangabeys from a single social group at TaïNP (0.0093% versus 0.016% sequence divergence, respectively). For the two African green monkey populations and the olive baboons, intra-group strain divergence was low (0.0004% and 0.0016%, respectively), though intra-species strain divergence for African green monkeys was almost as high as the divergence observed between the two most divergent human strains (0.009% versus 0.0093%). Although we do not have data to show ongoing NHP-to-human transmission (which is likely also influenced by a paucity of genomes from human-infecting strains), it is probable that the ancestor of *TPE* crossed species barriers more than once since neither human-nor NHP-infecting *TPE* strains form host species-specific clades. Moreover, the overall *TPE* clade exhibits a star-like phylogenetic branching pattern with low bootstrap support for short basal branches. This suggests a rapid initial radiation of *TPE* across humans and NHPs.

### Simian strains show the same genomic structure as human *TPE* strains

Using pooled segment genome sequencing (17) consisting of long-range PCR amplifications and NGS, we were able to determine the complete genome sequence (average depth of coverage: 169x) of the baboon strain LMNP-1 (ID: 40M5160407) collected in 2007 (13). The LMNP-1 genome showed the same structure as the complete genomes of three published human-infecting *TPE* strains Gauthier, CDC-2, Samoa D (25) and the simian strain Fribourg-Blanc (17). The genome of the LMNP-1 baboon strain (GenBank: CP021113) was more similar to the genome of human *TPE* Gauthier strain (NC_016843.1) than to simian isolate Fribourg-Blanc (NC_021179.1), showing differences at 266 and 325 chromosomal positions, respectively. Most differences between these genomes were single nucleotide substitutions or small indels. The LMNP-1 and Gauthier strains exhibited the same number of the 24-bp repeats in the *TP_0470* gene (n=25) and Gauthier had only one 60-bp repeat more than the simian strain LMNP-1 in the *arp* gene (LMNP-1 n=9 vs. Gauthier n=10). All 60-bp repeats in the *arp* gene of the LMNP-1 genome were of Type II and were identical to other *TPE* strains previously described (26). The *tprK* gene of the LMNP-1 isolate only had three variable regions, V5-V7, when compared to other *TPE* strains. In addition to differences in *TP_0433*, *TP_0470*, and *tprK* genes, relatively large indels were determined in *TPEGAU_0136* (33-nt long deletion; specific for strains Gauthier and Samoa D), in *TPFB_0548* (42-nt long deletion; specific for strain Fribourg-Blanc), in *TPEGAU_0858* (79-nt long deletion; specific for strain Gauthier), in the intergenic region (IGR) between *TPEGAU_0628* and *TPEGAU_0629* (302-nt long deletion; specific for strain Gauthier), and in IGR between *TPFB_0696* and *TPFB_0697* (430-nt long insertion; specific for strain Fribourg-Blanc); the length of other sequence differences ranged between 1-15 nts.

The structure of RNA operons of the LMNP-1 isolate (coordinates 231,180-236,139; 279,584-284,533; according to *TPE* strain Gauthier: NC_016843.1) was similar to strains Gauthier, CDC-2, and Fribourg-Blanc, but different to what was found in the strains Samoa D, Samoa F, and CDC-1 (27). The strain LMNP-1 sequence of 16S-5S-23S was identical in both operons and the 23S rRNA sequences were identical to other *TPE* strains except for strain Fribourg-Blanc (having a single nucleotide difference at position 458). Importantly, we have not found any mutations associated with macrolide resistance (e.g. A2058G, A2059G) (28, 29).

When the two NHP-infecting *TPE* strains, Fribourg-Blanc and LMNP-1, were compared to the closest human-pathogenic *TPE* strains CDC-2 and Gauthier, respectively, only 7.2% and 9.1% of all coding sequences (77 and 97 coding sequences out of 1065) were found to encode for amino acid substitutions. Comparisons of simian and human *TPE* genomes therefore suggest limited functional divergence. Since the Fribourg-Blanc strain causes sustainable infection when experimentally inoculated into human skin (30), it seems plausible that at least the LMNP-1 simian *TPE* strain has a zoonotic potential.

### A One Health approach is required to achieve sustainable yaws eradication

While reemerging yaws in humans cannot exclusively be explained by the existence of a natural reservoir in African NHPs (there are no NHPs in the Pacific region), there is compelling evidence that transmission routes between humans and NHPs exist. Since *TP* is mainly transmitted through direct contact, hunting infected NHPs (bushmeat) must be considered as a major risk for NHP-to-human infection (31). Pet monkeys may also function as a *Treponema*-reservoir for humans (32). Furthermore, it has been suggested that treponemes may be transmitted by flies (33-36). In this context it may seem surprising that Tanzania, where NHPs are infected with potentially zoonotic *TPE* strains (13), has not reported human yaws for decades. We note here this could either point towards very ineffective or no zoonotic transmission or clinical underreporting in humans. While mathematical modeling of human yaws-eradication suggests this goal can be achieved (37), our study provides clear evidence for the existence of *pertenue*-strains in NHPs, which points to the importance of a post-eradication surveillance program in human populations. We suggest holistic approaches taking into account the circulation of the bacterium in NHP populations, i.e. a One Health approach (38), will be needed to achieve sustainable eradication of human yaws post-2020.

## Materials and Methods

### DNA amplification, capture, and sequencing

Three different approaches were used to amplify and sequence the whole genomes of the simian *TP* strains. The method(s) depended on availability of DNA extracts and materials available in each laboratory. All West African *TP* isolates (TaïNP-1 and 2, BFP-1 and -2, NKNP-1 and -2) and the East African LMNP-2 strain were whole genome sequenced using in-solution and microarray based capture techniques, whereas the LMNP-1 strain was sequenced by microarray capture and long-range PCR. Long-range PCR generated high genome coverage in multiple repeat and paralogous regions where hybridization capture enrichment approaches proved less effective.

### In silico analysis

We applied EAGER (39), a comprehensive pipeline for read pre-processing, mapping, variant identification, and genome reconstruction on all sequenced samples. Each of the steps performed using EAGER is described briefly below.

#### Read preprocessing of sequenced genome samples

The sequenced products for all samples were paired-end reads with a varying number of overlapping nucleotides between corresponding forward and reverse reads. Several pre-processing steps were performed, including adapter clipping, merging of corresponding paired-end reads in the overlapping regions, and finally quality trimming of the resulting reads.

#### Mapping assembly

After adapter clipping, merging, and quality trimming, reads for all samples were mapped using Fribourg-Blanc (GenBank: NC_021179.1). All reads (merged and unmerged) were treated as single-end reads and mapped using the BWA-MEM algorithm (40) with default parameters. After mapping the Genome Analysis Toolkit (GATK) was used to generate a mapping assembly for each strain that had at least 80% coverage of the Fribourg-Blanc genome with a minimum of 3 reads. For this procedure, the UnifiedGenotyper module of GATK was applied to call reference bases and variants from the mapping. A reference base was called if the genotype quality of the call was at least 30 and the position was covered by at least three reads. A variant position (SNP) was called if the following criteria were met: i) the position was covered by at least three reads; ii) the genotype quality of the call was at least 30, and iii) the minimum SNP allele frequency was 90%. If the requirements for a variant call were not fulfilled, the reference base was called only if at least 3 reads confirmed it and the quality threshold was reached. If neither of the requirements for a reference base call nor the requirements for a variant call were met, the character ‘N’ was inserted at the respective position. Draft genome sequences were generated using the tool VCF2Genome of the EAGER pipeline.

#### SNP effect analysis

The genetic effect of each of the SNPs occurring in at least one strain was analyzed using SnpEff (40). We used an annotation database of *T. p. pallidum* (Fribourg-Blanc with RefSeq acc. ID NC_021179.1) built from the genomic annotation (the respective gff file was also retrieved from NCBI). These annotations include protein-coding genes as well as non-coding RNAs and pseudogenes. The up-/downstream region size parameter for reporting SNPs that are located upstream or downstream of protein-coding genes was set to 100 nt. For all other parameters default values were applied. Results were used to compile a table providing information on the genetic effect for each occurring SNP.

#### Longe range PCR-based genome assembly of the LMNP-1 TP genome

The Illumina sequencing reads obtained from the 4 distinct pools (sequenced as 4 different samples) were separately assembled *de novo* using SeqMan NGen v4.1.0 software (DNASTAR, Madison, WI, USA). A total of 99, 81, 62, and 138 contigs (obtained for pools 1, 2, 3, and 4, respectively) were aligned to the corresponding sequences of the reference Fribourg-Blanc genome (17) (GenBank: NC_021179.1) using Lasergene software (DNASTAR, Madison, WI, USA). In addition, Illumina sequencing reads were mapped to the Fribourg-Blanc genome and processed as described above. All gaps in the genome sequence and discrepancies between contig sequences and reference-guided consensus were resolved using Sanger sequencing. Altogether, 20 genomic regions of the baboon isolate genome from East Africa were amplified and Sanger sequenced. Overlapping pools were joined to obtain a complete genome sequence of the baboon isolate. The sequences of genes containing tandem repeats, i.e. *arp* (*TP_0433*) and *TP_0470* genes, were also resolved using Sanger sequencing. The number of tandem repeats in these genes was estimated using gel electrophoresis. Gene *tprK* (*TP_0897*) showed intra-strain variability and therefore nucleotides in variable regions were replaced with ‘ N’s in the complete genome sequence. In addition, the G/C-homopolymeric stretches revealed intra-strain variability throughout the genome. The prevailing number of G/Cs in these regions was used in the final genome sequence.

#### Genome identification, annotation, and classification

Both protein-coding genes and genes for noncoding RNA were annotated in the genome sequence of East African baboon isolate LMNP-1 based on the annotation of previously published *TPE* strain Gauthier (GenBank: NC_016843.1). Lasergene software was used for strain Gauthier orthologous gene alignment and recalculation of gene coordinates to East African baboon isolate LMNP-1. A 150-bp gene size limit was applied. Genes were tagged with the *TPE40M5*-prefix and the locus tag numbering corresponds to the tag numbering of orthologous genes annotated in the *TPE* strain Gauthier genome.

#### Processing of further human TPA and TPE samples from Arora et al. (2016)

Some samples of the extensive study conducted by Arora et al. (1) have been included for the phylogenetic analysis (see Table S3). Here, we only used those which achieved 80% coverage of the Fribourg-Blanc genome with a minimum of 3 reads, and which were Nichols-like (i.e. within the Nichols clade) or *TPE* samples. All raw reads were processed as described above.

#### Processing of published genomes

To apply the EAGER analysis pipeline to the complete genomic sequences already available in GenBank, we generated artificial reads using the tool Genome2Reads (also part of the EAGER software). Genome2Reads uses a tiling approach with an offset of 1 to artificially generate reads of 100nt reads, resulting in an average coverage of 100x. We applied the same mapping, SNP calling and genome reconstruction procedure as for the sequenced samples to obtain consistent and comparable results for phylogeny reconstruction.

#### Phylogenetic analysis

MEGA6 was used to generate maximum parsimony trees. Bootstrap values were inferred from 100 replicates. The analysis involved 28 nucleotide sequences, 8 NHP and 20 human strains. All positions with less than 85% site coverage were eliminated; in other words, fewer than 15% alignment gaps, missing data, and ambiguous bases were allowed at any position. We computed a phylogenetic tree using all 2,332 informative positions (**Fig. S1 A**) and another phylogenetic tree using the same alignment from which all positions from putative recombinant genes were removed (1), resulting in 2,063 informative positions (**Fig. S1 B**). Both trees were nearly identical and differed only in branch lengths, which can be attributed to the different number of informative positions.

## Acknowledgments

JFG was supported by an NSF Graduate Research Fellowship (DGE-1142336), the Canadian Institutes of Health Research’s Strategic Training Initiative in Health Research’s Systems Biology Training Program, an NSERC Vanier Canada Graduate Scholarship (CGS), and a long-term Research Grant from the German Academic Exchange Service (DAAD-91525837-57048249). HMD was supported by Max Planck Institute for Evolutionary Anthropology and Gent University. JK was supported by the Max Planck Society and the European Research Council (ERC) starting grant APGREID. KIB was supported by a Social Science and Humanities Research Council of Canada postdoctoral fellowship. The PSGS sequencing was supported by the Grant Agency of the Czech Republic to DS and MS (GA17-25455S, GJ17-25589Y). This project was supported by the German Research Foundation through grants FHL (LE1135/2), SK (KN1097/3-1 and KN1097/4-1), and CR (RO3055/2-1). This study was also supported by the AMIDEX project (No. ANR-11-IDEX-0001-02), funded by the 'Investissements d’Avenir' French Government program, managed by the French National Research Agency (ANR) and the Fondation Méditerranée Infection. All raw read files have been deposited in NCBI as part of the BioProject PRJNA343706. Please see supplementary acknowledgments for additional information.

